# Widespread mitochondrial DNA haplotypes of *Pteroptyx* spp. across present–day river systems in Southeast Asia and their historical dispersal through ancient river networks

**DOI:** 10.1101/2023.04.24.538207

**Authors:** Shawn Cheng, Nur H. Ali Akaram, Mohd A. Faidi, Tan Sek Aun, Subha Bhassu, Mohd N. Mat Isa

## Abstract

Estuarine fireflies from the genus *Pteroptyx* are widely distributed in Southeast Asia and famous for their nightly displays of bioluminescence by adult fireflies congregating or lekking on mangrove trees. *Pteroptyx* fireflies also offer insights into the history of the region, as their distribution in many of the isolated rivers they now inhabit are likely a product of dispersal via palaeorivers that formed in Southeast Asia during the Pleistocene. Here, we report the presence of widespread cytochrome oxidase subunit I (*cox1*) haplotypes among populations of *Pteroptyx* spp. in estuaries throughout Southeast Asia and suggest possible dispersal routes for these haplotypes vis-à-vis the Siam and Malacca River systems. Separately, reconstruction of the haplotype tree from the *cox1* gene indicated that the ancestors of *Pteroptyx asymmetria, Pteroptyx bearni, Pteroptyx malaccae, Pteroptyx tener*, and *Pteroptyx valida* either had a Thai, Bornean, or Peninsular Malaysian origin. Previous reconstructions of the phylogenetic and network trees of *Pteroptyx* sp. did not consider the presence of identical mitochondrial DNA haplotypes in their datasets, and the role palaeorivers had on their dispersal. The perspectives reported here aim to guide future taxonomic, phylogenetic and phylogeographic work on *Pteroptyx* fireflies in Southeast Asia.

## INTRODUCTION

Populations of *Pteroptyx* fireflies inhabit estuaries in Southeast Asia where they participate in spectacular displays of synchronised flashing on mangrove trees all year round (Ballantyne and McLean, 1970; Case, 1980; Lloyd et al., 1989). The pulsed flashing of *Pteroptyx* fireflies is attributed to male fireflies which have the unique ability to modulate their flashes, and which enables them to identify conspecifics and communicate with members of the opposite sex. Mass gatherings of synchronously and near-synchronously flashing populations of *Pteroptyx* spp. are a tourist attraction in Southeast Asia and an important contributor to the local economy (Cheng et al., 2021).

The life cycle of *Pteroptyx* fireflies is closely tied to the estuarine ecosystem. Females feed on the flowers of mangrove trees and mate with males on mangrove trees (Lloyd et al., 1989). Then they fly inland to lay their eggs amongst vegetation found in the intertidal zone. The larvae of *Pteroptyx* fireflies are semiaquatic and live in the intertidal zone where they hunt for estuarine snails for food (Nallakumar, 2003; Thancharoen et al., 2007). Larvae navigate the semiaquatic environment in the intertidal zone with the help of an organ called the pygopodia located at its hindmost segment which is used to attach itself to substrates, for motility and grooming (Archangelsky and Branham, 1998; Fu et al., 2005a).

*Pteroptyx* spp. often live in sympatry wherever they are found. Lloyd et al. (1989) reported the co-occurrence of *Pteroptyx malaccae* and *Pteroptyx valida* in mangroves in Tha Chin River, a tributary of Chao Phraya River in Thailand. Although *P. valida* was sometimes found alone in canals further away from the river, all localities where estuarine fireflies were found were always under tidal influence (Lloyd et al., 1989). Examination of published literature also reported the co-occurrence of *Pteroptyx tener, P. valida* and *P. malaccae* in Selangor River in Peninsular Malaysia; *Pteroptyx asymmetria, P. tener* and *P. valida* in Sepetang River, also in Peninsular Malaysia; and *P. malaccae* and *P. tener* in Tapi River in Thailand (see Table 1).

Reconstructions of the phylogenetic relationships of populations of *P. tener*, have in the past, consistently recovered two clades, one comprising Peninsular Malaysian samples along the west coast of Peninsular Malaysia, and Peninsular Malaysia populations from the east coast and a conspecific from Borneo, in another clade (Cheng et al., 2020; Jusoh et al., 2020). The pattern of Bornean conspecifics clustering with east coast sequences was also observed in *Pteroptyx bearni*, and to an extent, the clade comprising *Pteroptyx malaccae* and *Pteroptyx balingiana* (Jusoh et al., 2020). Jusoh et al. (2020) also considered the “…spatio-genetic structure of populations within *P. tener, P. bearni, P. malaccae* and *P. balingiana*…” (sic.) to be an indicator that cryptic species complexes existed.

Although Jusoh et al. (2019) alluded to the possibility that palaeoriver systems may have had a role in the dispersal of *Pteroptyx tener* populations in Peninsular Malaysia, their ultimate view was that the phylogeographic information imprinted onto their phylogenetic tree was a result of cryptic or incipient speciation (Jusoh et al., 2020). There has also been failure to appreciate the presence of identical mitochondrial DNA (mtDNA) haplotypes inhabiting a wide range of present-day tributaries or rivers in Southeast Asia in the datasets of Jusoh et al. (2019, 2020) and Yaakop et al. (2023). Climate change driven range displacements was previously shown to have contributed to the dispersal of *P. tener* into different river systems in Peninsular Malaysia (Cheng et al., 2020). Seeing that ever since, there have been missed opportunities to address the cause behind the unusual distribution of *Pteroptyx* fireflies in Southeast Asia, we reassessed the datasets of Sartsanga (2018), Cheng et al. (2020), and Jusoh et al. (2020) to provide some unifying insights or explanations into the presence of identical DNA haplotypes in otherwise isolated estuarine river systems in Southeast Asia.

## MATERIALS AND METHODS

### Data collection and analysis

*Cytochrome oxidase subunit I* (*cox1*) gene sequences of *Pteroptyx asymmetria, Pteroptyx bearni, Pteroptyx malaccae, P. tener*, and *Pteroptyx valida* were downloaded from GenBank with the Batch Entrez tool (NCBI, USA). Datasets for each species were aligned with BioEdit 7.2.5 using the ClustalW algorithm (Hall,1999) and were used to identify identical haplotypes of *Pteroptyx* spp. in PopArt (Bandelt, 1999).

### Phylogenetic tree construction and analysis

We reconstructed the haplotype tree of *Pteroptyx* spp. with *cox1* haplotype sequences of *P. asymmetria, P. bearni, P. malaccae, P. tener*, and *P. valida*. We used the following species to root our phylogenetic sequences: *Colophotia brevis* (KY572911), *Colophotia praeusta (*KY572909), Luciolinae sp. (KY572913), *Pteroptyx galbina* (KY572914), and *Pteroptyx testacea* (KY572919). The unaligned FASTA sequence file containing all sequences used in this report can be found in Supporting Information S1.

We used RAxML-HPC2 on XSEDE (8.2.12) (Randomized Axelerated Maximum Likelihood), a phylogenetic analysis tool on CIPRES (Miller et al., 2010) to reconstruct our phylogenetic tree and performed 1000 bootstrap replicates to calculate the robustness of our estimated tree. Output trees were visualised in IcyTree, a browser-based phylogenetic tree viewer (Vaughan, 2017) and annotated with information on species, haplotype identities and localities where the samples were collected from.

### Haplotype distribution on present-day river systems superimposed on Pleistocene Sea level maps

We superimposed a Pleistocene Sea level map over a present-day map of Southeast Asian rivers. Then, we mapped the distribution of all five *Pteroptyx* spp. using data from Sartsanga et al. (2018), Cheng et al. (2020), and Jusoh et al. (2020). Following this, we assessed the connectivity of populations inhabiting present-day river systems to ancient rivers that formed in Southeast Asia at the point when sea levels were 120 m below the present level (BPL).

## RESULTS

### General patterns in the cox1 haplotype tree of co-occurring Pteroptyx species from Southeast Asia

Figure 1 shows the reconstruction of our *cox1* haplotype tree comprising *Pteroptyx asymmetria* (Clade I), *Pteroptyx tener* (Clade II), *Pteroptyx bearni* (clade III), *Pteroptyx valida* (Clade IV), and *Pteroptyx malaccae* (Clade V). *P. asymmetria* was recovered at an ancestral position with Thai populations occupying a basal position and a mixture of Thai and Peninsular Malaysian sequences from the west coast, at a derived position (Figure 1). This was followed by a second subclade comprising Thai sequences of *P. asymmetria* at a basal position and Peninsular Malaysian sequences from the west coast at a derived position (Figure 1). A similar pattern of Thai sequences resolving at an ancestral position and Peninsular Malaysian firefly populations from the west coast appearing at a derived position, was also observed in *P. tener* (Clade II, subclade B) and *P. valida* (Clade IV) (Figure 1). Separately, we recovered Bornean sequences occupying a basal position followed by Peninsular Malaysian samples from the east coast at a derived position in *P. tener* (Clade II, subclade A) and *P. bearni* (Clade III) (Figure 1). We also detected several haplotypes that found their way into different river systems in Southeast Asia. In *P. asymmetria*, for example, *cox1* haplotype ‘A’ was found inhabiting river systems in Thailand and the west coast of Peninsular Malaysia (Figure 1, Table 1). Separately, haplotype B of *P. bearni* was found to have dispersed into two different locations along the east coast of Peninsular Malaysia, in Chukai River (Terengganu) and Lebam River (Johor) (Table 1).

**Figure 1.**
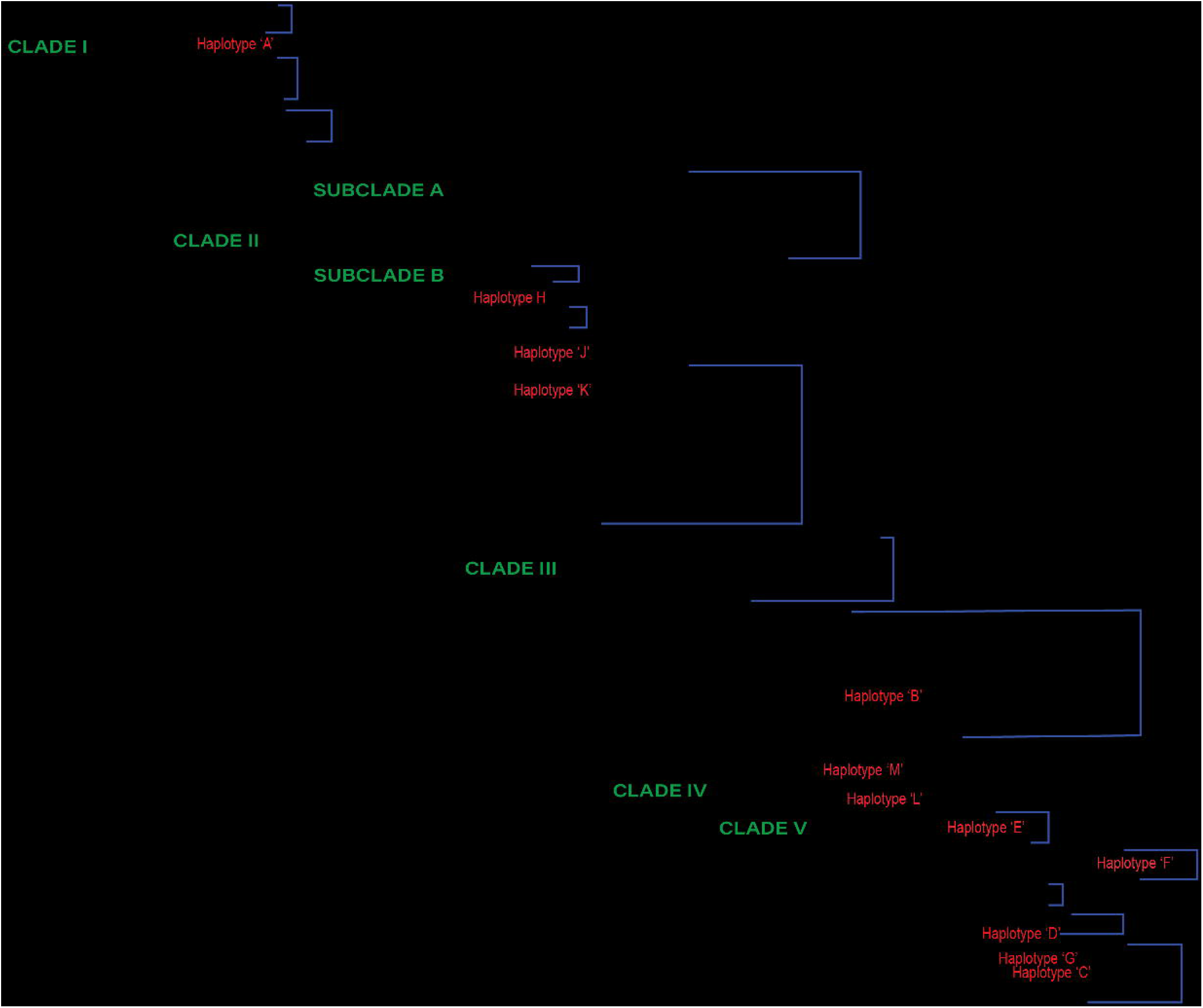
Maximum likelihood haplotype tree estimated with RAxML. Only branch support values above 50% are shown on the tree.

### Pteroptyx tener

A single Bornean sequence of *Pteroptyx tener* (KY572989) gave rise to the east coast populations of the species with both recovering as a strongly supported clade, as opposed to *P. tener* samples from the west coast of Peninsular Malaysia and Thailand which resolved into another clade. We also detected three widespread *P. tener* haplotypes, that is, haplotypes ‘H’, ‘J’, and ‘K’ (Figure 1). Haplotype ‘H’ was found in Tapi River in southern Thailand, and Sepetang and Selangor Rivers on the west coast of Peninsular Malaysia (Figure 1, Table 1). Haplotype ‘J’ meanwhile was found in Rembau, Linggi and Sepetang Rivers on the west coast of Peninsular Malaysia. Separately, Haplotype ‘K’ was found in Rembau and Linggi Rivers in Negeri Sembilan (Figure 1). Both these rivers however merge into a larger tributary prior to draining into the Straits of Malacca.

### Pteroptyx bearni

*Pteroptyx bearni* was recovered at an ancestral position compared to *P. valida* and *P. malaccae* with the node leading to the clade showing high support (Figure 1). The clade comprised sequences from Suai and Raan Rivers in Sarawak (Borneo); Lebam River on the east coast of Johor, Chukai River in Terengganu, and Pahang Tua River on the east coast of Pahang (all from Peninsular Malaysia). Sequences from Suai and Raan Rivers (Borneo) occupied a basal position in Clade III followed by a clade comprising east coast samples that were descendants of the Bornean samples. The node leading to *P. bearni* individuals from the east coast had a high bootstrap value (Figure 1). Lastly, we detected one widespread haplotype, that is, Haplotype ‘B’, that was found in both Chukai River (Terengganu) and Lebam River (Johor), in the east coast of Peninsula Malaysia.

### Pteroptyx valida

Clade IV consisted of a basal sequence from Samut Songkhram (southwest of Bangkok) and sequences from southern Thailand, Peninsular Malaysia, and Borneo, all occupying a derived position (Figure 1). Among the derived sequences were two haplotypes, that is, haplotype ‘L’ which was found in Merbok, Sepetang and Pak Bara Rivers (along the western coast of the Malay Peninsula) and haplotype ‘M’ which dispersed into estuaries in the Gulf of Thailand, the west coast of Peninsular Malaysia and Borneo (Figure 1).

### Pteroptyx malaccae

Clade V comprised of a basal sequence from the west coast of Peninsular Malaysia and sequences from Thailand and Peninsular Malaysia (west and east coast populations) at a derived position. We found five haplotypes occupying multiple river systems in Southeast Asia e.g., haplotype ‘D’ in Chukai and Pahang Tua Rivers (east coast, Peninsular Malaysia) and haplotype ‘E’ in Raya and Rembau Rivers (west coast, Peninsular Malaysia) (Figure 1, Table 1). In the *cox1* haplotype tree, two identical sequences of *P. malaccae*, that is MH431847-48 resolved on a vertical line. These sequences shared a close relationship with *P. malaccae* sequences from the east coast of Peninsular Malaysia including haplotype ‘D’ which was found in Chukai and Pahang Tua Rivers (Figure 1, Table 1).

## DISCUSSION

### Palaeoriver systems in Southeast Asia and their effect on firefly dispersal

Land bridges that formed during the Pleistocene in the Sunda shelf promoted the dispersal of terrestrial organisms throughout Southeast Asia (Husson et al. 2019). Apart from the formation of land bridges, the Pleistocene also saw the formation of palaeorivers in the region which aided the dispersal of freshwater fishes and snakes (Inger and Chin, 1962; Geyh et. al., 1979; Karns et al., 2000; Voris, 2000). The estuarine firefly, *Pteroptyx tener*, also shared a similar dispersal history (Cheng et al., 2020) through movement of mangrove swamps past present-day shorelines. Physical evidence of the movement of mangrove swamps past present-day shorelines is supported by discovery of spore-pollen assemblages along the east coast of Peninsular Malaysia. Biswas (1973) reported the presence of *Pandanus-Barringtonia* transitional swamp, nipa palm tidal swamp and *Rhizophora-Sonneratia* mangrove swamp assemblages hundreds of feet below the sea floor in offshore gas exploration fields, here.

An alternative but untested suggestion was that the dispersal of one *Pteroptyx* species, that is, *Pteroptyx tener*, could have been facilitated by the ability of its larvae to ride river or ocean water currents to reach new areas (Yaakop et al., 2023). Although fully aquatic larvae can exchange gases under water (Jeng et al., 2003), there is no evidence of the presence of gills in the larvae of *P. tener*. Taxonomic descriptions of the larva of its nearest relatives, that is, *Pteroptyx valida* and *Pteroptyx maipo* found the absence of gills along the sides of their body, that would normally be found in aquatic fireflies (Ballantyne and Menayah, 2002; Ballantyne et al., 2011).

Dispersal during the Pleistocene did not affect *P. tener* alone, as the data presented in this report shows that it has affected its congeners as well. This was firstly by virtue of them occurring in sympatry wherever they have been found and secondly, by the discovery of widespread, identical haplotypes of *Pteroptyx asymmetria, Pteroptyx bearni*, Pteroptyx malaccae, *P. tener*, and *Pteroptyx valida* throughout their distribution in Southeast Asia (Figure 1, Table 1). Widespread haplotypes occupying different river systems in Southeast Asia provide an early indication of historical/past migration events.

*Pteroptyx bearni* haplotype ‘B’, for example, was found in Chukai and Lebam Rivers, both of which may have been connected to the Siam River system in the South China Sea during the Pleistocene (Figure 2). Separately, *P. tener* haplotype ‘J’, by virtue of being found in Sepetang, Rembau and Linggi Rivers, likely dispersed via the Malacca River System during the Pleistocene (Figure 2). Although these rivers were not among the major palaeorivers draining into the Malacca River System, Voris (2000) suggested that there could have been no fewer than four large rivers contributing to the flowage between the Malay Peninsula and Sumatra. However, complexities arise when describing dispersal in the haplotypes of other taxa within the *Pteroptyx*. Haplotype ‘A’ of *P. asymmetria* and Haplotype ‘H’ of *P. tener*, were found in mangroves from two disjunct localities, that is, the Gulf of Thailand and the west coast of Peninsular Malaysia (Figure 1, Table 1).

**Figure 2.**
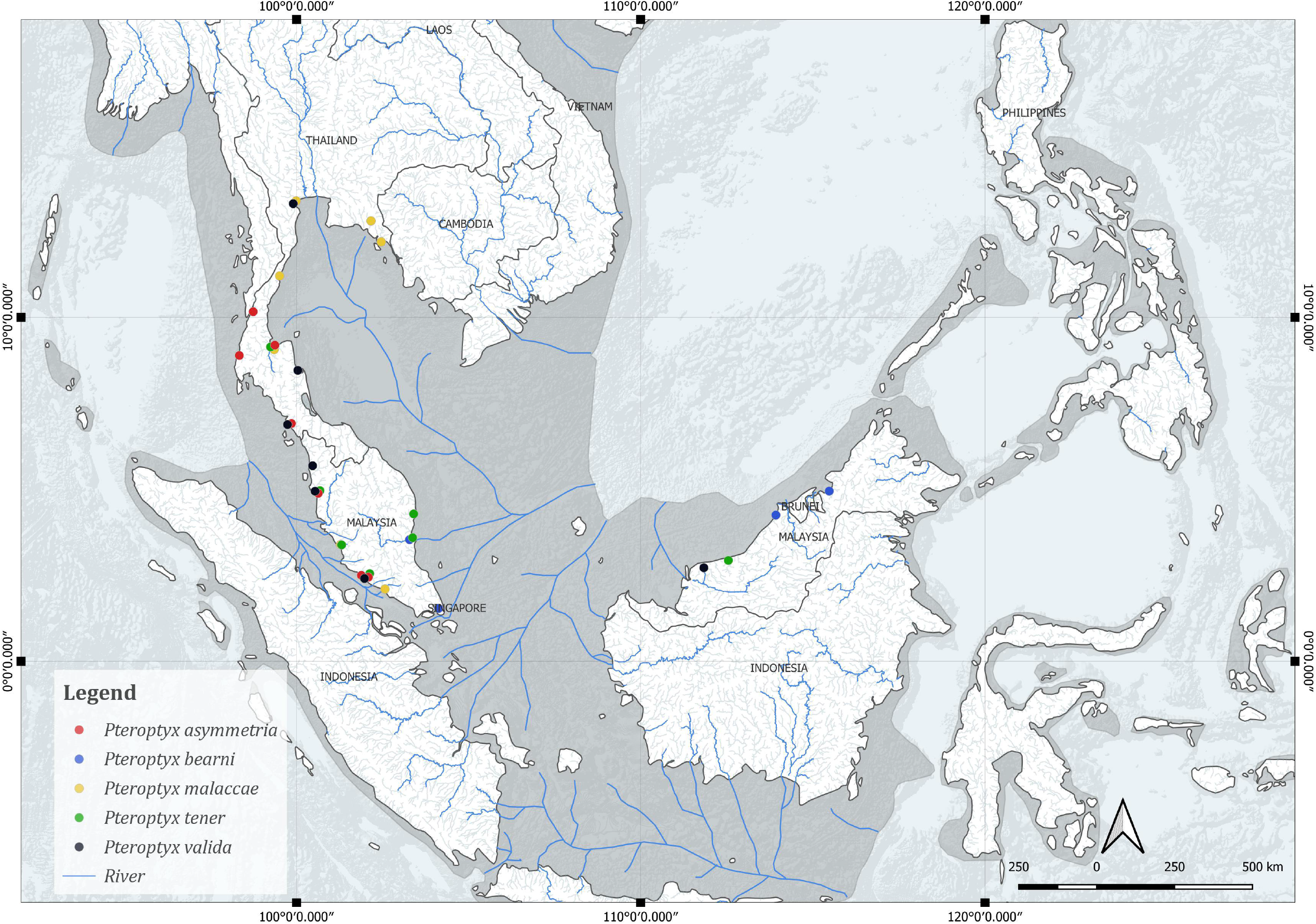
Pleistocene rivers at the 120-meter level superimposed onto a contemporary map showing Southeast Asian River systems. Locations in coloured circles indicate distribution of Pteroptyx spp, as reported in the literature.

### Identification of ancestral populations of Pteroptyx spp. in Southeast Asia

In two *Pteroptyx* clades, that is, *Pteroptyx asymmetria* and *Pteroptyx tener* (Clade II, subclade B), ancestral populations were Thai in origin. Descendants of these populations eventually found their way to the west coast of Peninsular Malaysia (Figure 1). Dispersal to the west coast of Peninsular Malaysia could have occurred when Sumatra’s Sungai Kampar, which ran north through the Singapore Straits and Johore River, joined branches of the Siam River System when sea levels were 120 m below present level (Voris, 2000).

Connectivity between the Siam River system and the Malacca River system meanwhile may have been facilitated by the formation of a large freshwater estuary between the Malay Peninsula and Sumatra when sea levels rose to 20 m BPL during the Pleistocene (Voris, 2000). This freshwater estuary could have been a refugia for populations of *Pteroptyx* spp. from the Siam River System that later dispersed into the Malacca River System when sea levels dropped again during the Pleistocene. In the past 250,000 years, there were between two and six events when sea levels fluctuated from 10 m BPL to 120 m BPL (Voris, 2000) and which may have provided opportunities for their dispersal to the west coast of the Malay Peninsula. To test this hypothesis, there is a need to collect and sequence adequate numbers of firefly samples from these locations. Following this, one would then need to identify dispersal scenarios between adjacent populations of these species and estimate the most probable scenario describing the history of these biogeographically disjunct populations of *Pteroptyx* spp. (see Cornuet et al., 2010, 2014; Cheng et al., 2020).

The non-uniform distribution of *Pteroptyx* haplotypes and species in Southeast Asia, however, could also be due to insufficient data. Until there is an exhaustive effort to sample *Pteroptyx* species and the sharing of genetic resources between firefly researchers in Southeast Asia, it may be premature to make further conclusions about their otherwise disjunct distribution in the region.

Sea level fluctuations during the Pleistocene which brought about cycles of connectivity and isolation between populations served as drivers of divergence/vicariance due to increases and decreases in the size of terrestrial habitats (Papadopoulou and Knowles, 2015; Husson et al., 2019). In the case of estuarine fireflies, the Pleistocene brought about relatively long periods of connectivity vis-à-vis palaeoriver systems which formed in the Sunda Shelf followed by prolonged periods of isolation/vicariance between the river systems in here (Voris, 2000). The Pleistocene is generally thought to be a period when active population differentiation occurred, and only when environmental conditions permitted their survival and divergence between intraspecific phylogeographic assemblages, the creation of new species (Avise and Walker, 1988).

## Conclusions

Climate change driven range displacements that occurred during the Pleistocene likely facilitated the dispersal of *Pteroptyx* spp. throughout Southeast Asia. Long periods of connectivity from the formation of palaeoriver systems during this epoch provided dispersal opportunities for *Pteroptyx* spp. while prolonged periods of isolation/vicariance between these river systems brought about speciation. Failure to appreciate the presence of widespread mtDNA haplotypes in our DNA datasets, and the role that palaeorivers had on dispersal in estuarine fireflies has clouded our ability to understand the processes that shaped their distribution in the entire region today. This is likely the first report detailing how estuarine insects dispersed with the help of palaeorivers during the Pleistocene. The perspectives detailed here should guide future taxonomic, phylogenetic and phylogeographic work on *Pteroptyx* fireflies in Southeast Asia.

## Supporting information

Table 1. cox1 haplotypes of Pteroptyx spp. and their distribution in Southeast Asia.

## ACKNOWLEDGEMENTS

We are indebted to Harold K. Voris from the Field Museum of Natural History (Chicago, USA) for permission to use and reproduce palaeoriver maps published by the Museum for this article. Finally, we thank the Director General of FRIM and the Director of the Biotechnology Division for resources provided to the first author to embark on this manuscript. All authors contributed equally to this article.

## SUPPORTING INFORMATION

**Table 1.** *cox1* haplotypes of *Pteroptyx* spp. and their distribution in Southeast Asia. Sequences from Sartsanga et al. (2018), Cheng et al. (2020), Jusoh et al. (2020). TH= Thailand; PM= Peninsular Malaysia; SRW= Sarawak in Borneo.

**Supporting Information S1**. FASTA sequences of unique mtDNA haplotypes and outgroup sequences used to estimate the maximum likelihood tree.

