## Supplementary material for "Widespread mitochondrial DNA haplotypes of *Pteroptyx* spp. across present–day river systems in Southeast Asia and their historical dispersal through ancient river networks": Table 1. cox1 haplotypes of Pteroptyx spp. and their distribution in Southeast Asia.

**Table 1.** *cox1* haplotypes of *Pteroptyx* spp. and their distribution in Southeast Asia. Sequences from Sartsanga et al. (2018), Cheng et al. (2020), Jusoh et al. (2020). TH= Thailand; PM= Peninsular Malaysia; SRW= Sarawak in Borneo.

| **Species** | **Haplotype Identity** | **Distribution** | **GenBank Accession Nos.** |
| --- | --- | --- | --- |
| *P. assymetria* | ‘A’ | Sepetang River (West PM)  Laun, Ranang (TH)  Takua Pa (TH)  Khlong Thom (TH)  Krabi (TH) | KY572923  MF624791  MF624796  MF624797  MF624801 |
| *P. bearni* | ‘B’ | Chukai River (East PM)  Lebam River (East PM) | KY572925,30,41-42,45-47,53  KY572938 |
| *P. malaccae* | ‘C’ | (TH)  Tapi River (TH) | MF624806  MF624807 |
|  | ‘D’ | Chukai (East PM)  Pahang Tua (East PM) | KY572963  KY572965 |
|  | ‘E’ | Raya River (West PM)  Rembau River (West PM) | KY572959-60  KY572961 |
|  | ‘F’ | Ban Nam Chiao (TH)  Kung Krabaen (TH) | MF624813  MF624816 |
|  | ‘G’ | Prachuap Kiri Khan (TH)  Tapi River (TH)  Samut Songkhram (TH) | MF624804  MF624805  MF624809-12,18,20-22 |
| *P. tener* | ‘H’ | Sepetang River (West PM)  Selangor River (West PM) | KY572969,71-72,77-78  MH431832  MH431864 |
|  | ‘K’ | Tapi River (TH)  Rembau River (West PM)  Linggi River (West PM) | MF624824,26,28  KY572973  MH431889 |
|  | ‘J’ | Sepetang River (West PM)  Rembau River (West PM)  Linggi River (West PM) | KY572970,79  KY573011, MH431839  KY572976,86  KY573010,27  MH431881 |
| *P. valida* | ‘L’ | Merbok River (West PM)  Sepetang River (West PM)  Satun (TH) | KY573050  KY573051  MF624830-31 |
|  | ‘M’ | Raya River (West PM)  Mukah River (SRW)  Muang (TH) | KY573046-47  KY573048  MF624829 |
